# Inconsistent short-term effects of enhanced structural complexity on soil microbial properties across German forests

**DOI:** 10.1101/2025.07.15.664741

**Authors:** Rike Schwarz, Nico Eisenhauer, Christian Ammer, Pia M. Bradler, Orsi Decker, Benjamin M. Delory, Peter Dietrich, Andreas Fichtner, Yuanyuan Huang, Ludwig Lettenmaier, Michael Junginger, Oliver Mitesser, Jörg Müller, Goddert von Oheimb, Kerstin Pierick, Michael Scherer-Lorenzen, Simone Cesarz

## Abstract

Structural and biotic homogenization can result from forestry practices that lack promotion of canopy gaps and deadwood. This can lead to biodiversity loss and impaired ecosystem functions. Enhancing structural complexity (ESC) has been proposed to counteract these effects, but its impact on soil properties remains insufficiently understood. Overall, we hypothesize that ESC enhances soil abiotic properties, their spatial variability, and microbial functioning, with effects modulated by environmental context and increasing over time. Data were collected from 148 patches (50 × 50 m) in eight beech forests across Germany. In half of the patches, structural complexity was enhanced by felling 30% of the basal area of living trees through two spatial patterns—aggregated (one large gap) and distributed (small gaps)—combined with leaving or removing deadwood (stumps, logs, and snags). The other half served as controls, representing typically managed, homogeneous production forests. Soil C:N, C%, and N% increased near deadwood. Soil microbial biomass and activity were significantly affected in three of eight forest sites, with effects ranging from −30% to +62%. Higher soil water content was associated with increased microbial biomass, and greater understorey biomass correlated with a lower microbial respiratory quotient. No temporal trends were observed over five years. Although soil properties showed resistance to structural interventions, site-specific effects underline the importance of soil moisture and the understorey vegetation for microbial functioning. Further research building on our results is needed to develop practical forest management strategies to clarify how structural complexity may support soil functioning and ecosystem resilience.

**Graphical abstract:** 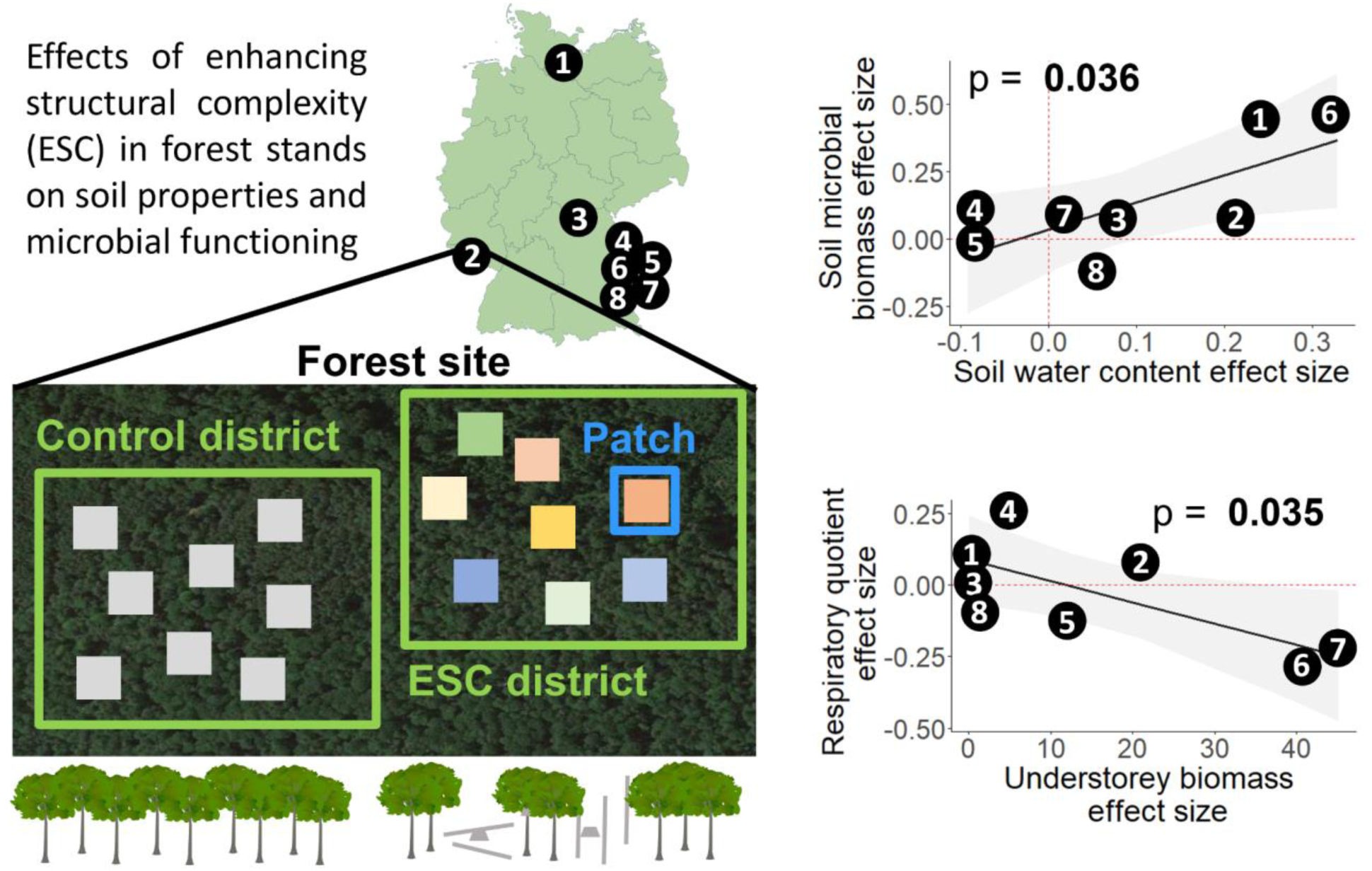

**Highlights:**

- Deadwood addition increases soil C%, N%, and the soil C:N ratio
- Enhanced structural complexity alters soil microbial properties in site-specific ways
- Soil water content changes are linked to shifts in microbial biomass
- Understorey biomass changes are linked to shifts in the respiratory quotient

## 1. Introduction

Homogenization of landscapes is a serious human-made threat to biodiversity (McGill et al., 2015). In production forests, intensive silvicultural management can lead to landscape homogenization by creating stands missing early-successional stages, with usually small canopy gaps and little deadwood (Aszalós et al., 2022; Paletto et al., 2014). This reduction of habitat diversity can cause biodiversity loss (Newbold et al., 2015; Rousseau et al., 2019) and impair ecosystem functioning (Hooper et al., 2012; Soliveres et al., 2016). This has far-reaching consequences for human well-being (Cardinale et al., 2012; Wall et al., 2010). Thus, it is essential to identify sustainable approaches for managing forests for timber production while promoting biodiversity and maintaining multiple ecosystem functions (Messier et al., 2022; Topanotti et al., 2023). Structural complexity of forests has received growing attention (Beugnon et al., 2024; Ray et al., 2023), alongside timber harvesting methods that aim to retain key structural elements, such as standing live trees and deadwood (*e.g.*, Churchland et al., 2021; Lewandowski et al., 2019). Here, we consider the expansion of canopy gaps and the presence of deadwood in a forest as enhancing its structural complexity (Keeton, 2006; Müller et al., 2023). In gaps, more light and water reach the forest floor (Gálhidy et al., 2006; Latif & Blackburn, 2010; Ritter et al., 2005), while deadwood can alter soil properties (Błońska et al., 2017; Moghimian et al., 2020). However, there is a knowledge gap regarding the effect of forest structure and complexity on belowground organisms that drive important ecosystem functions.

Current estimates suggest that ∼59% of all species on Earth inhabit the soil (Anthony et al., 2023), and they play a crucial role in the mineralization of organic matter, nutrient cycling, and carbon (C) sequestration (Bardgett & van der Putten, 2014; van der Heijden et al., 2008). Soil basal respiration, microbial biomass, and the respiratory quotient are useful indicators for soil ecosystem functions (Cesarz et al., 2022; Eisenhauer et al., 2010). For example, higher soil microbial biomass correlates with higher wood mass loss (Gottschall et al., 2019), and higher soil microbial activity correlates with soil C sequestration (Lange et al., 2015).

Soil microbial community diversity, biomass, and activity are sensitive to nutrient availability, soil pH, and moisture (Fierer & Jackson, 2006; Mulder et al., 2005; Serna-Chavez et al., 2013). Enhancing a forest’s structural complexity through added canopy gaps and deadwood can influence these variables, and it likely also affects soil biodiversity and functioning. For example, forest gaps can result in higher understorey biomass and diversity (Gálhidy et al., 2006; Mueller et al., 2016), which in turn can enhance root exudates and increase soil microbial biomass (Chen et al., 2019; Eisenhauer et al., 2017; Lange et al., 2015). Deadwood, in turn, provides an important substrate for fungi and other wood-decomposing organisms (Dove & Keeton, 2015; Dyson et al., 2024). Depending on tree species, type (stump/log), and decay stage, deadwood can enhance soil water content, soil organic C, total nitrogen (N), enzymatic activity, and soil respiration, while contrasting effects on soil pH have been recorded (Błońska et al., 2017; Moghimian et al., 2020; Perreault et al., 2020; Piaszczyk et al., 2019; Wambsganss et al., 2017). Taken together, canopy gaps and deadwood can lead to higher variability of physicochemical soil properties due to the patchy impact of deadwood as well as the heterogeneity of the regenerating vegetation in canopy gaps (Moghimian et al., 2020; Perreault et al., 2020; Ritter et al., 2005). As a result, a more heterogeneous habitat for soil organisms can increase their diversity (Curd et al., 2018; Eisenhauer, 2016). More diverse communities are often linked to increased functioning (Bell et al., 2005; Colombo et al., 2016). Therefore, we expect that increased structural complexity leads to changes in both abiotic and biotic soil properties.

Different environmental contexts in a forest might modulate the effect of increased structural complexity on soil organisms and their activity. Here, we refer to the concept of mechanistic context dependency (Catford et al., 2022), where the relationship between variables (e.g., enhanced structural complexity and biotic soil properties) is influenced by interaction effects with other factors (e.g. abiotic soil properties or the understorey vegetation). Since soil water and nutrient availability are strong predictors of soil microbial properties (Serna-Chavez et al., 2013), the potential changes in these abiotic properties at a site may affect the influence of other factors on soil microbial properties (Cesarz et al., 2022). For instance, in experimental forest stands, tree diversity and identity were reported to have a greater impact on soil microbial properties in drier and nutrient-poor soils than in more moist and nutrient-rich soils (Cesarz et al., 2022; Lu & Scheu, 2021). Another example is that the effect of structural complexity on the understorey may vary with soil conditions (Leuschner & Ellenberg, 2017), potentially influencing the effect on soil microbial properties (Eisenhauer et al., 2010; Lange et al., 2015). Thus, we expect that the strongest microbial responses may occur at sites where structural complexity induces the largest changes in soil abiotic conditions or understorey vegetation, such as in relatively dry or nutrient-poor forests, highlighting the importance of environmental context in shaping belowground responses to structural interventions.

While mechanistic context dependency highlights the variability of biodiversity-ecosystem functioning relationships across different spatial contexts, effects might also change over time. Experimental research indicates that the impact of biodiversity on ecosystem functioning becomes more pronounced over time (Bongers et al., 2021; Guerrero-Ramírez et al., 2017; Reich et al., 2012), but there is a lack of time series data on forest soils investigating such relationships. As the effects of plant species richness on belowground properties (Ravenek et al., 2014; Strecker et al., 2016) and the influence of deadwood on soil develop gradually (Moghimian et al., 2020; Perreault et al., 2020), the impact of structural complexity on soil microbial properties may also change over time.

This study aims to address these research gaps by investigating the effects of enhanced structural complexity of forest stands on abiotic soil properties, soil microbial activity, biomass, and the respiratory quotient in different environmental contexts in eight forests in Germany. Time series data are available for five years for one of these forests. We hypothesize that (H1a) canopy gaps and deadwood enhance soil water content, pH, and nutrient levels and (H1b) increase their spatial variability. As a consequence, we expect (H2) a general increase in soil microbial activity, biomass, and a decrease of the respiratory quotient, indicating an increase of the efficiency of the microorganisms, in forests with greater structural complexity. Thirdly (H3), we expect this effect to be mechanistically context-dependent and more pronounced in forest sites where enhanced structural complexity has a stronger effect on soil water content, soil nutrient availability, and the understorey vegetation. Finally (H4), we expect the influence of enhanced structural complexity on soil microbial activity, biomass, and the respiratory quotient to increase over the first five years.

## 2. Methods

### 2.1. Study design

This study was conducted in the BETA-FOR experiment (Müller et al., 2023). To investigate the hypotheses in different environmental contexts, eight deciduous - primarily beech-dominated - forest sites in Germany were selected, and study patches were established (Müller et al., 2023). One site is located in the forest of the University Würzburg (U03), five sites are in and close to the Bavarian Forest National Park (B04, B05, B06, B07, P08), one site is close to Saarbrücken (S10), and one close to Lübeck (L11; Figure 1A). The forest sites differ in their environmental context (Table 1; Figure 2). Each forest site consists of a control district and a treatment district (Figure 1B). In the treatment district, the structural complexity was enhanced (Enhancement of structural complexity = ESC districts) by creating canopy gaps and deadwood through different silvicultural interventions. These treatment districts will be referred to as ESC districts. The interventions in the ESC districts were always applied to 30 % of the stand basal area of living trees of the 50 x 50 m patch. The tree removal was conducted in two spatial variants: aggregated and distributed. In the aggregated variant, all trees were removed centrally, creating one large contiguous gap (30 m diameter). In contrast, in the distributed variant, trees were removed uniformly throughout the whole patch, resulting in small-scale canopy openings, while the same volume of timber as in the aggregated variant was removed. Further, four deadwood treatments were crossed with the two spatial variants (aggregated vs distributed): stumps, logs, snags, or logs and snags remaining (Figure 1C). To create snags, trees were cut below the crown with a harvester to retain standing deadwood. Forest site U03, an add-on and a long-term site, included six additional patches accommodating three more deadwood treatments to allow investigation of a higher level of spatial heterogeneity (though not the focus of this study). Across all forest sites, deadwood treatments resulted in an average deadwood volume equivalent to 45 m^3^ deadwood hectare^-1^ (± 29 m^3^). Patches in the control district were not treated and thus represent a normally thinned forest or a currently unmanaged former commercial forest (in the national park). Within each district (10 - 20 ha), there are nine patches, each measuring 50 x 50 meters. In the ESC districts, one of these nine patches remained untreated to increase within-district heterogeneity. Given that our study focuses on the effects of treated (ESC) versus untreated (control) conditions on soil properties and microbial functioning, we excluded the untreated patch within the ESC district from all analyses to avoid biased results. The contiguous control and treatment districts are in close proximity and have similar geology, soil types, and tree species composition. Thus, in total, 70 ESC patches ((7 forests * 8 patches) + 14 (the eighth forest with 14 patches) = 70) and 78 control patches ((7 forests * 9 patches) + 15 (the eighth forest and all control patches) = 78) were included in our analysis, resulting in 148 patches overall.

**Figure 1:**
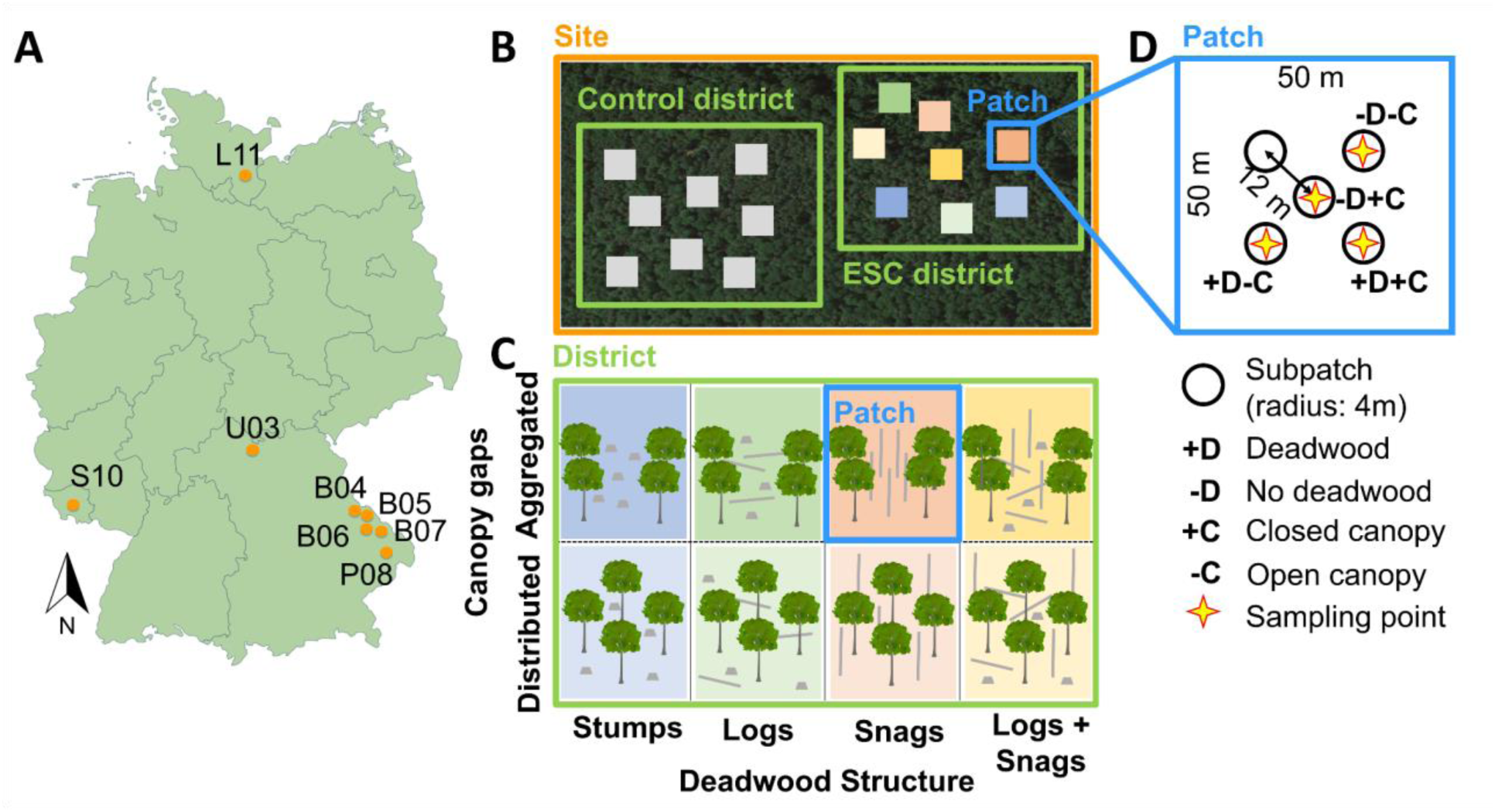
Locations of the study sites across Germany (A), schematic representation of a site with the ESC (enhancement of structural complexity) and the control district (B), illustration of the individual treatments within an ESC district along two axes, i.e., canopy gaps in aggregated and distributed configuration and different deadwood structures (C), and schematic representation of the sampling method (D). We captured within-patch heterogeneity by sampling across combinations of canopy cover (open vs. closed) and deadwood presence. Ideally, one sample was taken from each of the following: (1) deadwood under open canopy, (2) no deadwood under open canopy, (3) deadwood under closed canopy, and (4) no deadwood under closed canopy. Sampling began at the patch center, from which three subpatches were selected to best represent all condition combinations. Panel (D) shows an example distribution of sampling points and conditions within a patch.

**Figure 2:**
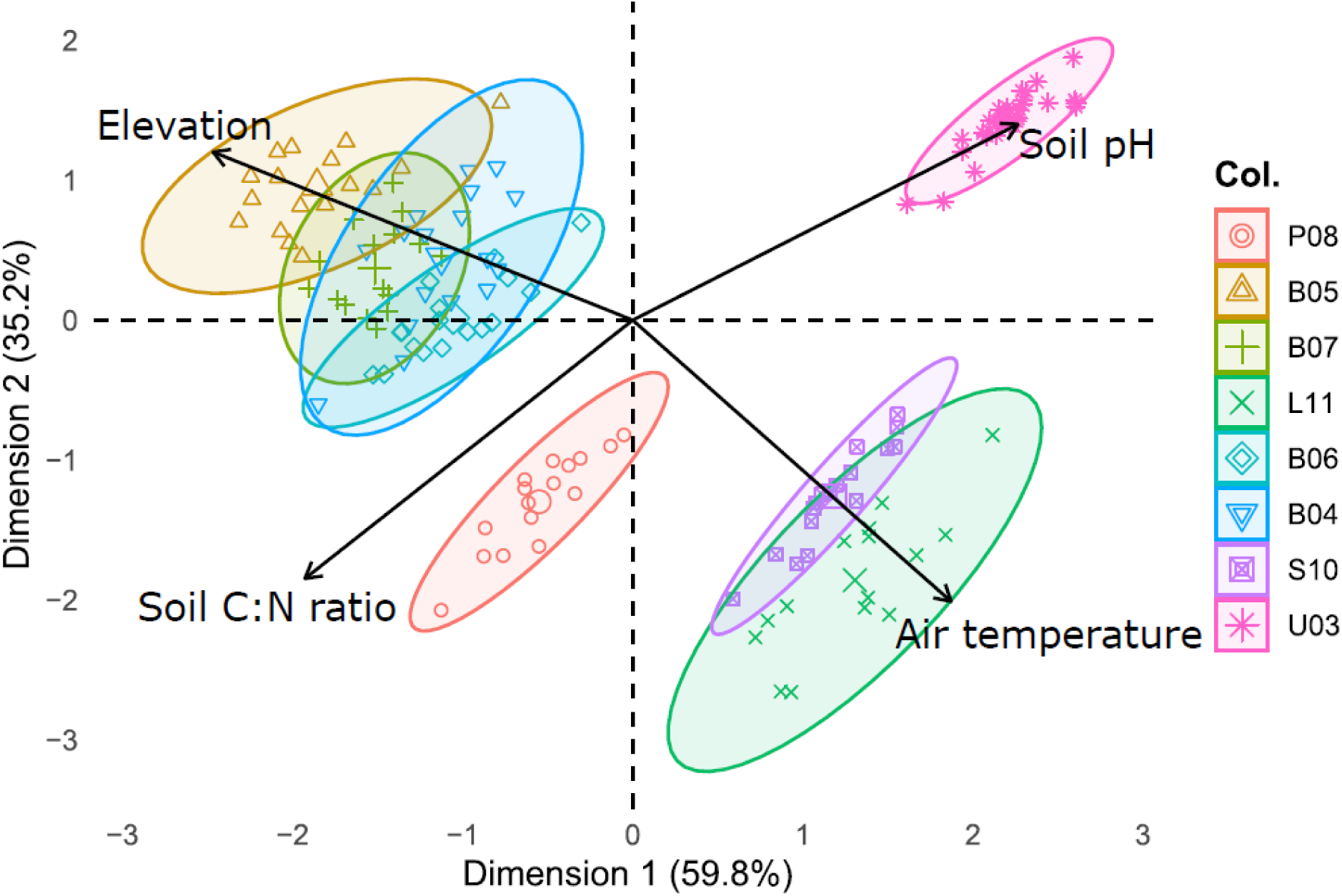
Principal component biplot of the environmental context of patches of this study comprising elevation, soil pH, air temperature, and soil carbon-to-nitrogen (C:N) ratio. Data was standardized, points are color-coded for different forest sites and each ellipse shows a forest site with 95% confidence intervals.

**Table 1:** Environmental context of the eight forest sites (ordered by pH), including the year the ESC districts were established (Year of experimental intervention). For temperature (Temp.), soil C:N ratio, soil pH (measured in CaCl_2_ solution), and soil water content, the mean of the control patches was used to characterize each forest site. Temperature refers to the average at 2 m height in 2023, measured at the patch center. Geology/lithology, soil type, and vegetation type each represent the dominant category within the respective forest site (ESC and control patches).

| Forest site | Elevation [m] | Temp. [°C] | C:N ratio | pH | Year of experimental intervention | Geology/Lithology | Soil type | Vegetation type |
| --- | --- | --- | --- | --- | --- | --- | --- | --- |
| P08 | 496 | 11.3 | 22.6 | 2.7 | 2015/16 | Carboniferous/ granite | Weakly podzolic Cambisol | *Luzulo-Fagetum* |
| B05 | 1080 | 8.2 | 19.3 | 2.7 | 2015/16 | Carboniferous/ granite | Weakly podzolic Cambisol | *Luzulo-Fagetum* |
| B07 | 928 | 9.0 | 19.4 | 2.8 | 2015/16 | Carboniferous/ granite | Weakly podzolic Cambisol | *Luzulo-Fagetum* |
| L11 | 41 | 13.4 | 19.1 | 3.0 | 2016/17 | Young moraine/ basal till | Weakly podzolic stagnic Cambisol | *Galio-Fagetum* |
| B06 | 829 | 9.8 | 19.4 | 3.0 | 2015/16 | Carboniferous/ granite, gneiss | Cambisol | *Luzulo-Fagetum* |
| B04 | 888 | 9.4 | 17.8 | 3.0 | 2015/16 | Carboniferous/ gneiss | Weakly podzolic Cambisol | *Luzulo-Fagetum* |
| S10 | 296 | 13.2 | 19.3 | 3.9 | 2015/16 | Upper carboniferous/ silt- and mudstone | Stagnic Cambisol | *Galio-Fagetum* |
| U03 | 311 | 10.6 | 13.6 | 5.7 | 2018/19 | Muschelkalk/<br>limestone | Chromic<br>Cambisol | *Hordelymo-*<br>*Fagetum* |

### 2.2. Soil sampling and analyses

Four soil cores (diameter 5 cm, depth 10 cm) per patch were taken within a 12 m radius around the center of the patch after roughly removing the litter layer. The four sampling points were selected to represent the heterogeneity of the patch with the canopy gaps and deadwood axes. This resulted in samples taken next to deadwood vs. no deadwood and open vs. closed canopy (Figure 1D). This was done in October 2023 on all sites. U03 was additionally sampled each autumn since 2018 to study the ESC effect over time. Samples were constantly cooled and sieved (2 mm) before further processing. Soil C, N, and pH were measured on the sampling point level (4 per patch); microbial biomass, basal respiration, and soil water content were measured from samples pooled per patch.

Prior to microbial measurements, samples were kept for three days at +20°C to acclimate the soil microbial community to a consistent and standardized temperature. We used an automated O_2_-microcompensation apparatus (Scheu, 1992) to assess soil basal respiration and microbial biomass. Basal respiration represents the mean oxygen consumption per hour (μl O_2_ h^-1^ g soil dw^-1^) without any substrate addition. This indicates the active portion of the soil microbial community at the time and soil condition of sampling. For this, the mean oxygen consumption between the hours 10 - 20 was taken. Microbial biomass C was determined through substrate-induced respiration, which measures the respiratory response of microorganisms to added glucose and water (Anderson & Domsch, 1978). An aqueous solution containing 8 mg of glucose per gram of dry soil was added. The lowest substrate-induced respiration observed over three consecutive hours within the first 10 hours was taken as the maximum initial respiratory response (MIRR), taking place before microbial growth started. Microbial biomass (μg C g^-1^ soil dw) was calculated as 38 x MIRR, based on calibration against fumigation methods (Beck et al., 1997). This method measures the full potential of the living microbes that can use glucose. The respiratory quotient is calculated by dividing the basal respiration by the microbial biomass. It determines the ratio of built-up C to investment in respiration. If the respiratory quotient is low, substrate use is efficient, as more microbial biomass can be built up with less respiration. We use the respiratory quotient as an indicator of substrate use efficiency and refer to it as such.

Soil water content was determined by weighing the fresh sample, drying it at 75°C for three days, and reweighing it (IAEA, 2008; Sünnemann et al., 2021). As this measurement reflects the soil water content at the moment of sampling, it is only suitable to compare different patches that were sampled under the same weather conditions. Accordingly, ESC and control districts of the same forest site were always sampled within a short period of time (*i.e.,* within a maximum of four days), while different forest sites were sampled up to three weeks apart. To avoid, for example, a heavy rain event at one forest site influencing our results, soil water content is only used for comparison within a forest site and not between forest sites. To measure soil pH, 10 g of air-dried soil was mixed with 25 ml of 0.01 M CaCl_2_ solution, shaken, and allowed to sit for 1 h. The pH was then recorded using a pH meter (Orion Star A211, Thermo Scientific, MA) (Bönisch et al., 2024; FAO, 2021). For determination of C and N content, soil samples were dried at 30°C for 72 h, ground, and then transferred into tin capsules (20 mg each). The analysis was conducted using dry combustion with a Vario EL cube IR elemental analyzer (Dietrich et al., 2021; Farina et al., 1991). The C and N content were provided as the percentage of the element’s mass relative to the sample mass, and the C:N ratio was calculated from these values. Soil water content was measured at the patch level from a pooled soil sample, while all other abiotic soil variables were recorded for every sampling point per patch (n = 4).

### 2.4. Further environmental variables

During the vegetation period of 2023, five hemispheric photographs were taken on all patches to measure the canopy openness. One photograph was captured at the center of each patch, while the others were taken between 12 and 20 m apart from the center in a cross-shaped pattern in all four cardinal directions. No people or equipment appeared in the images. The camera used was a Nikon D7200, equipped with a Sigma 4.5 mm F2.8 EX DC Circular Fisheye HSM lens, mounted on a tripod at a height of 1 m and oriented directly upwards. The diffuse site factor (DIFFSF), representing the proportion of diffuse solar radiation penetrating the canopy, was calculated from these images using “Segmentation” software v1.0.0.1 (developed by the Chair of Photogrammetry, TUD, in 2006) and “Hemisphere Tool v1.1 beta” (February 2018, developed by the same chair). For further analyses, an average DIFFSF per patch was used and referred to as canopy openness.

The deadwood volume was assessed on all patches. On the ESC patches, all deadwood that was experimentally created (standing, lying, stumps, habitat trees) was recorded. On the control patches, only naturally dead standing trees and fresh stumps from thinning were considered. For standing deadwood, a taper of 1 mm per decimeter was assumed. The volume of crowns and lying deadwood was calculated using the formula: 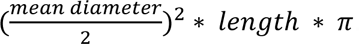. Deadwood with a diameter under 7 cm, including snags and logs, was excluded from documentation. Thus, the value corresponds to the coarse woody debris present. For simplicity, we will refer to deadwood throughout the manuscript.

Air temperatures at 2 m height were measured with one EL-USB2 (Lascar Electronics Ltd., UK) data logger per patch. The loggers were installed in TX COVER (Technoline, Germany) logger shields, which were mounted on wooden poles at the patch centers. Air temperatures were measured every 30 minutes. Data gaps, resulting from logger malfunctions, were imputed from information of loggers from the same district using the R package ‘MissForest’ (Stekhoven & Bühlmann, 2012). The temperature time series were aggregated to mean temperatures of the year 2023 for each site.

Surveys and aboveground biomass harvest of the understorey were conducted in the vegetation period of 2023 on five subpatches (4 m radius each) per patch. During the vegetation surveys, all herbaceous and woody species up to the height of 1 m were recorded. For understorey biomass harvest, in each subpatch, app. 0.5 m^2^ (70 x 70 cm) of aboveground understorey vegetation was harvested in a location that was representative of the patch and subpatch in terms of species biomass and richness. The biomass was oven-dried (60°C for 72 h) and weighed. Biomass values were summed per patch and standardized to 1 m^2^.

### 2.6. Statistical analysis

All statistical analyses were conducted in R v.4.2.2 (R Core Team, 2020). To test for differences in environmental conditions (soil pH, air temperature at 2 m, soil C:N ratio, elevation) among forest sites, we performed a principal component analysis (PCA) using the function *prcomp* from the package ‘stats’ (R Core Team, 2022).

To test whether the forest interventions that increase light availability and deadwood occurrence affect abiotic variables (soil water content %, soil pH, soil C:N ratio, soil C%, and soil N%), we used both continuous and categorical explanatory variables (H1a). As continuous variables, we used canopy openness (the proportion of visible sky) to represent light and deadwood volume (in m³). Categorical variables included the spatial pattern of tree removal (control vs. distributed vs. aggregated) and the presence of deadwood within 30 cm of the sampling point (yes or no). Importantly, deadwood volume (continuously; overall deadwood volume per patch) and deadwood occurrence (categorically; spatially explicit presence of deadwood next to sampling point) represent distinct concepts. Deadwood occurrence was recorded for each sampling point whereas all other explanatory variables were used at the patch level. We fitted one model for each abiotic soil variable separately, using tree removal variant, canopy openness, deadwood volume, and deadwood occurrence as fixed effects without including interaction terms. Deadwood volume was log-transformed due to its non-linear relationship with many variables (Müller et al., 2010). For the patch-level data (soil water content %), forest site was included as a random effect, while for data on sampling point level (soil C%, N%, soil C:N ratio, and soil pH), patch nested within forest site was used as a random effect.

To test if our interventions lead to higher heterogeneity in our response variables (H1b), we calculated the coefficient of variation (CV) for non-pooled samples per patch (n = 4). We fitted a linear mixed-effects model with the CV as the response variable, district (control vs. ESC) as fixed factor, and forest site as random effect. Given that soil water content was only determined at the patch level (and not at the higher spatial resolution), we did not test the effect of deadwood occurrence on soil water content (H1a) and also not the effect of district (control vs. ESC) on the CV of soil water content (H1b).

To test whether ESC affected the microbial functions (soil basal respiration, soil microbial biomass, and respiratory quotient) and if the effects on the three biotic soil properties differ between forest sites (H2), we fitted linear mixed-effect models with district and forest site, and their interaction as fixed effects, respectively. The devices on which basal respiration and microbial biomass were measured was always included as a random effect.

To test if greater increases in soil abiotic and understorey properties lead to greater increases in soil biotic properties (H3), we calculated a mean per forest site and district combination (*e.g.,* U03 control and U03 ESC) for soil basal respiration, soil microbial biomass, respiratory quotient, soil water content, soil C:N ratio, soil C%, understorey plant species richness and aboveground biomass. We calculated the relative difference between ESC and control district for every forest site and variable, hereafter referred to as effect size. As we cannot form pairs of control and ESC patches, this had to be done at the district level. We then tested the relationship between the effect sizes of abiotic/ understorey variables and effect sizes of biotic variables. To get an overview of forest site-specific effects of ESC on the abiotic soil variables, we fitted models with district, forest site, and their interaction as explanatory variables.

To test if the effect of ESC on soil properties increased over the first five years of the experiment (H4), we analyzed data from 2019-2023 from U03. Since the samples from 2018 were taken only two weeks after ESC establishment, we excluded this year’s data to avoid any disturbance effects. We fitted linear mixed-effects models with district (categorical variable) and year (continuous variable), as well as their interaction as fixed effects and patch and measurement device as random effects. Further, we calculated the effect size of ESC on soil basal respiration, soil microbial biomass, and the respiratory quotient for each year (relative difference between control and ESC district, respectively) and fitted linear models with year as a continuous variable as the explanatory variable and the effect size as the response variable.

Throughout the study, we fitted beta regression models for percentage data (beta distributed errors), and for non-percentage data, we used models with normally distributed errors. Linear models were fitted using the package ‘stats’ (R Core Team, 2022), linear mixed-effects models were fitted using the ‘lme4’ package (Bates et al., 2015), and beta regression models were fitted using the package ‘glmmTMB’ (Brooks et al., 2017). The function *anova* from the package ‘stats’ (R Core Team, 2022) was used to produce p-values. For mixed-effects models, p-values for fixed effects were computed using the ‘lmerTest’ package (Kuznetsova et al., 2017), which provides Satterthwaite’s degrees of freedom approximation. Assumptions of the models were visually checked using the R package ‘performance’ (Lüdecke et al., 2021). If necessary, variables were log-transformed to improve model fit. This was done for pH, C:N CV, C% CV, and N% CV for H1 analyses and for basal respiration and the respiratory quotient for H2 analyses. To assess the significant interactions, post-hoc tests were conducted using the ‘emmeans’ package (Lenth, 2021), and compact letter displays were derived using the package ‘multcomp’ (Hothorn et al., 2008). For all figures, predicted values (marginal effects) were computed using the function *ggpredict* from the package ‘ggeffects’ (Lüdecke, 2018). All graphs were created with the ‘ggplot2’ package (Wickham, 2016). The estimated marginal means from the post-hoc tests were used to calculate the change of soil properties in percentage.

## 3. Results

### 3.1. Effects of canopy gaps and deadwood on physicochemical properties of soil (H1)

The spatial variant of tree removal, canopy openness, and deadwood volume per patch did not significantly affect soil water content, soil pH, soil C:N ratio, soil C%, and soil N% across all forest sites (Figure 3, Table S1 in supporting information). Deadwood occurrence near the sampling point did not change soil pH significantly (Figure 3G), but significantly increased the soil C:N ratio (+ 2%; Figure 3K), soil C% (+ 8%; Figure 3O), and soil N% (+ 6%; Figure 3S). Neither soil pH, nor the C:N ratio, nor C% or N% showed a higher CV in the ESC districts than in the control districts (Figure S1 and Table S2 in supporting information).

**Figure 3:**
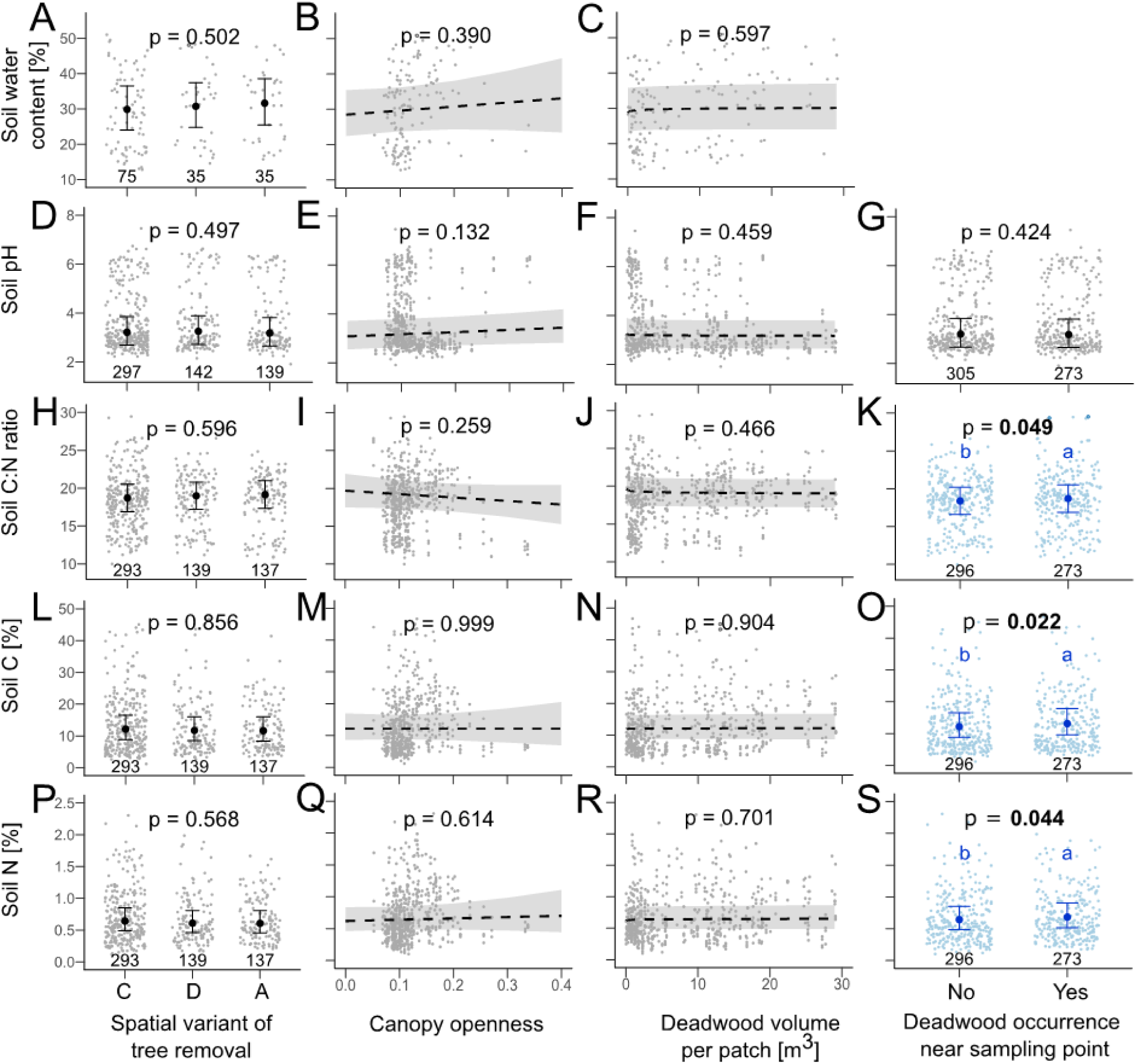
Soil water content (A - C), soil pH (D - G), soil C:N (H - K), soil C% (L - O), and soil N% (P - S) as affected by spatial variant of tree removal (left column; C= control, D= distributed, A= aggregated), canopy openness (second column), deadwood volume per patch (second column from right; m^3^), and deadwood occurrence near sampling point (right column). P-values are from beta regression models (soil water content, C%, N%) and linear mixed-effect models (soil pH, soil C:N ratio); significant results (p < 0.05) are bold and panels highlighted in blue. Model predictions are shown with 95% confidence intervals; dashed lines indicate non-significant effects, and raw data are shown in the background. Sample sizes (n) are indicated for categorical predictors.

### 3.2. Effect of ESC on soil respiration, microbial biomass, and the respiratory quotient (H2)

Across forest sites, soil basal respiration, soil microbial biomass, and the respiratory quotient did not differ between control and ESC districts (though district was marginally significant for microbial biomass, post-hoc tests did not show a significant difference between districts). However, the interaction between forest site and district was significant for basal respiration, indicating that the specific location is important (Figure 4, Table S3 in supporting information). Specifically, basal respiration was significantly higher in the ESC district in L11 (+ 62%) and lower in the ESC district in P08 (−30%; Figure 4A). The effect of the interaction between forest site and district on soil microbial biomass was marginally significant and showed a significant increase in the ESC district in B06 (+ 42%; Figure 4B). The respiratory quotient was not significantly different between control and ESC districts in any of the forest sites (Figure 4C).

**Figure 4:**
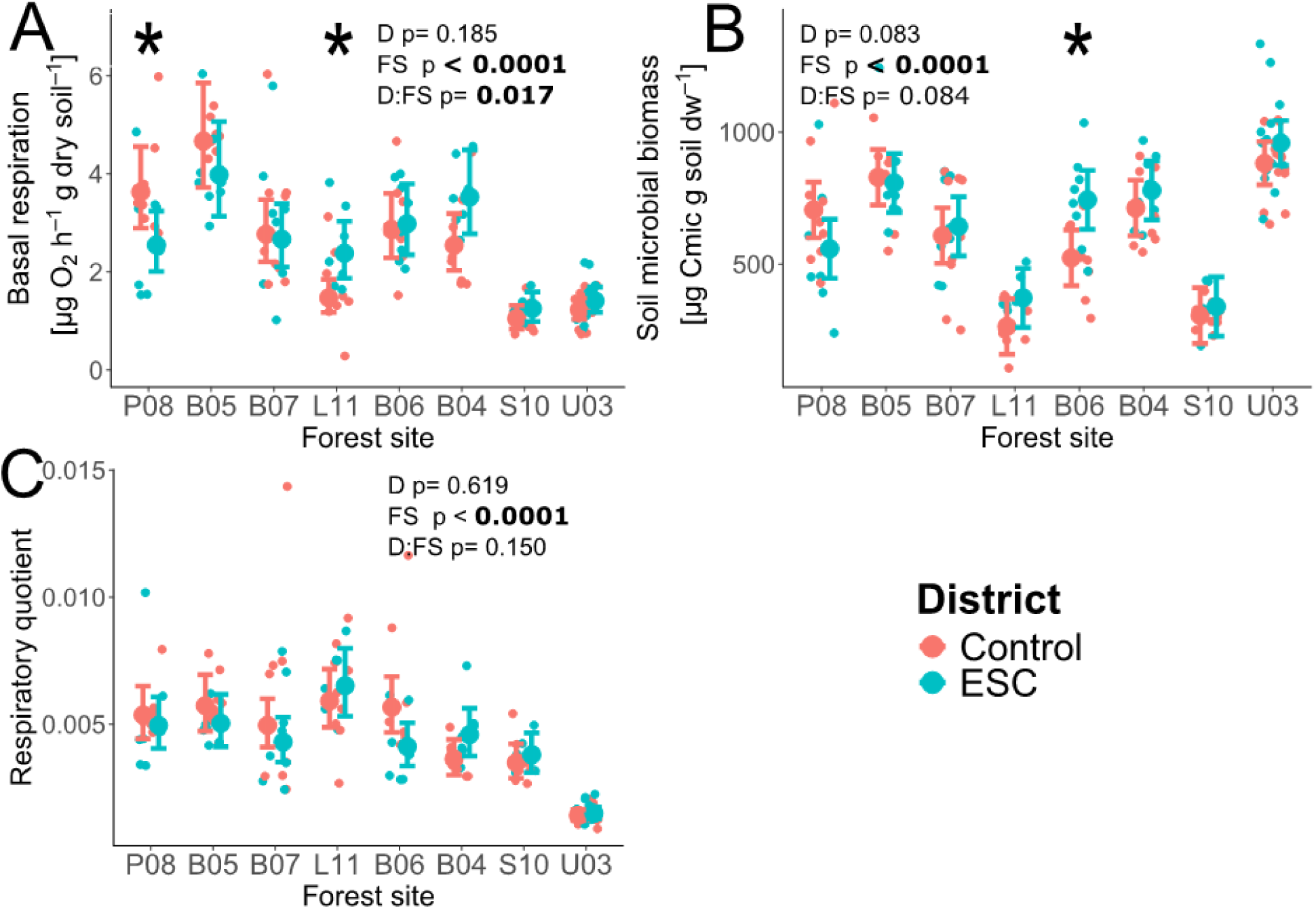
Soil basal respiration (A), soil microbial biomass (B), and soil respiratory quotient (C) in 2023 per forest site and district (control: red; enhanced structural complexity (ESC): blue). P-values are from linear mixed-effect models (D= District; FS= Forest site); significant values (p < 0.05) are bold. Asterisks indicate significant within-site differences from post-hoc tests following (marginally) significant FS × D interactions. Model predictions are shown with 95% confidence intervals (error bars); larger points indicate means, raw data per patch in the background. Forest sites are ordered by increasing soil pH.

### 3.3. Mechanistic context dependency of ESC effects (H3)

Here, we tested if the relative change between control and ESC in one variable (=effect size) is associated with a corresponding effect size of another variable for 15 combinations of a variable (abiotic soil properties and understorey vegetation properties) with a biotic microbial variable (Figure 5, Table S4 in supporting information). This could indicate a positive or negative correlation, depending on the trend of the data, or a lack of association if the changes in one effect size don’t appear to predict changes in the other. In two out of the 15 combinations, we found a significant relationship, and in one combination, a marginally significant effect (Figure 5). We observed a significant positive relationship between the change in soil microbial biomass and soil water content (Figure 5B). The effect size of ESC on the respiratory quotient was significantly negatively affected by the effect size of ESC on understorey biomass (Figure 5L) and marginally negatively affected by observed plant species richness (Figure 5O). An overview of ESC effects on abiotic soil properties can be found in Table S5 in the supporting information. Soil pH was not affected by ESC in any of the forest sites. Soil water content was significantly higher in the ESC districts of three forest sites (B06: + 35%; L11: + 23%; S10: + 26%). Soil C:N ratio was significantly increased by ESC in one forest site and decreased in another (B04: + 11%; P08: - 8%). Soil C% and N% were significantly lower in the ESC district of one forest site (B05: - 31% (C%); −29% (N%)).

**Figure 5:**
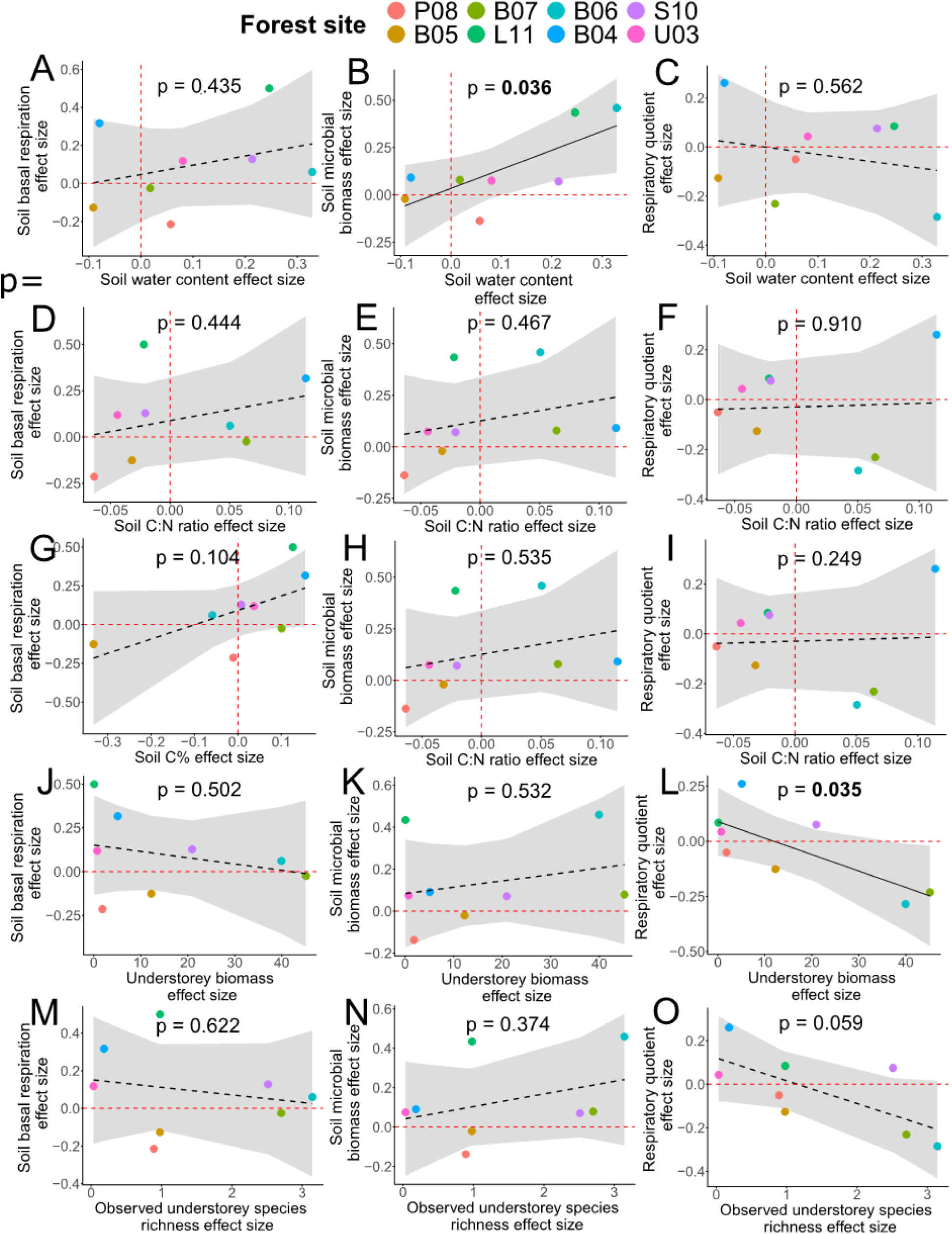
Relationships between the effect size of enhanced structural complexity (ESC) on soil basal respiration (left), soil microbial biomass (middle), and respiratory quotient (right) with the effect sizes of ESC on soil water content (A - C), C:N ratio (D - F), C% (G - I), understorey biomass (J - L), and observed understorey plant species richness (M - O). Black lines show model predictions with 95% confidence intervals; points show the raw data, color-coded by forest site. P-values are from linear models; significant values (p < 0.05) are bold. Significant relationships are shown with solid lines and non-significant ones with dashed lines. The red line indicates the zero baseline; values above it show an increasing effect and values below it a decreasing effect by ESC.

### 3.4. Temporal development of ESC effects over the first five years (H4)

ESC did not significantly influence soil basal respiration, soil microbial biomass, and the respiratory quotient across the five years of investigation in U03 (Figure 6, Table S6 in supporting information). Soil basal respiration and respiratory quotient decreased significantly with time, whereas soil microbial biomass was stable. No significant interaction of year and district was found for any of the three soil biotic properties. Year had no significant effect on the effect size (relative difference in mean values) between control and ESC districts for the response variables basal respiration, microbial biomass, and respiratory quotient in U03 (Figure S2 and Table S7 in supporting information).

**Figure 6:**
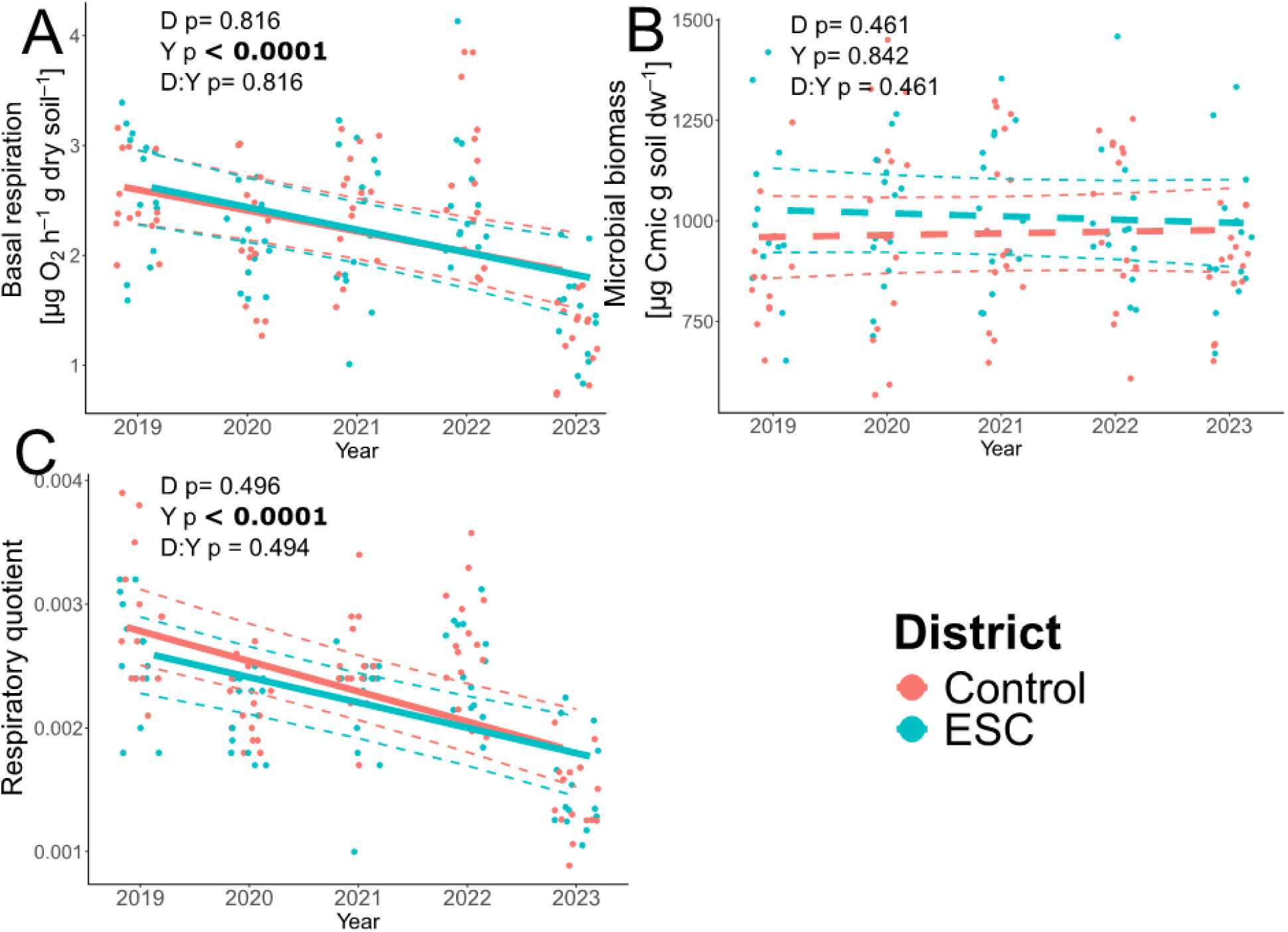
Soil basal respiration (A), soil microbial biomass (B), and soil respiratory quotient (C) per year and district (control: red; enhanced structural complexity (ESC): blue) from forest site U03. P-values are from linear mixed-effect models (D= District; Y= Year); significant values (p-value < 0.05) are bold. Lines represent model predictions with 95% confidence intervals, points show raw data per patch. Significant relationships are shown with solid lines, non-significant ones with dashed lines. Thick lines for basal respiration and the respiratory quotient are displayed in bold to illustrate overall temporal trends, despite the non significant D x Y interaction.

## 4. Discussion

Due to intensive management, production forests frequently become structurally homogenized on the landscape scale, leading to a loss of biodiversity and ecosystem functions. Here, we studied the effects of experimentally enhanced structural complexity (ESC) in forests on soil abiotic and biotic properties over space and time.

### 4.1. Effects of canopy gaps and deadwood on physicochemical properties of soil (H1)

Enhanced light availability and deadwood only partially influenced the physicochemical properties of the soil (H1). Soil pH and soil water content remained unchanged, while soil near deadwood showed an increase in C%, N%, and the C:N ratio. Even though several studies observed increased soil water content in canopy gaps within one to two years after gap creation (Gálhidy et al., 2006; Latif & Blackburn, 2010; Ritter et al., 2005), our findings align with a study on 3–6-year-old gaps that found no consistent increases (Lenk et al., 2024). These results suggest that increases in soil water content are only short-term and may be influenced by gap size and closure dynamics. In smaller gaps, like those in our distributed variant, rapid closing due to the crown plasticity of neighboring trees (Schröter et al., 2012) may lead to higher interception by the canopy, while in larger gaps, like our aggregated variant, increasing natural regeneration may take up the available soil water (Kovács et al., 2020; Ritter et al., 2005). Similarly, we also found no effect of increased deadwood volume on soil water content, likely because the mechanisms by which deadwood enhances soil moisture—through absorbing and storing water, and incorporating organic material into the soil—tend to intensify gradually over time with ongoing decomposition (Błońska et al., 2018; Edelmann et al., 2023; Piaszczyk et al., 2019). However, increased soil C% and N% near deadwood suggest some organic material has already been integrated into the soil, the effects remain weak and have not yet influenced soil water content. During early decomposition, deadwood can act as a net N source (Bantle et al., 2014). However, fungal N translocation from surrounding substrates into deadwood (Schimel & Hättenschwiler, 2007) may counteract this effect, which could explain why we observed only a weak increase in soil nitrogen around deadwood. The effects on soil C% and N% near deadwood, but not for deadwood volume on patch level, emphasize the small-scale effect of deadwood on soil, as shown in previous studies (Minnich et al., 2021; Moghimian et al., 2020; Stutz et al., 2017). This, in turn, underscores the crucial role of deadwood in fostering a heterogeneous soil habitat for microorganisms.

Soil pH was not altered by the presence of deadwood or open canopies. Even though literature is inconsistent about the effect of deadwood on soil pH, strong effects are often associated with highly decayed deadwood (Moghimian et al., 2020; Perreault et al., 2020, Stutz et al., 2017). For instance, Gonzalez-Polo et al. (2013) and Błońska et al. (2023) observed the most significant impact on soil properties when deadwood had reached advanced decay (stage V: no leaves, twigs, or bark, with a soft, powdery wood consistency), whereas the deadwood in our patches is better classified as medium decay stage (stage II: leaves absent, but twigs and bark present, with a solid wood consistency). Importantly, the development of deadwood into higher decay stages can take decades (Vrška et al., 2015). Contrary to previous studies (*e.g.* Perreault et al., 2020), variability (CV) of physicochemical soil properties did not increase in ESC patches. Even though this was not expected, it is consistent with the generally weak effect on the abiotic soil properties and may change as time progresses.

### 4.2. Effect of ESC on soil respiration, microbial biomass, and the respiratory quotient (H2)

Enhanced structural complexity did not generally increase soil basal respiration, microbial biomass, or decrease the respiratory quotient across the different forest sites (H2). Nevertheless, ESC increased basal respiration in one of the sites and decreased it in another, while soil microbial biomass increased in ESC patches of a third site. These inconsistent effects suggest a mechanistic context dependency (H3). The previously discussed absence of strong general effects of ESC on the physicochemical soil properties is in line with the absence of an overall effect on biotic soil properties as these two sets of properties are functionally connected (Fierer & Jackson, 2006; Serna-Chavez et al., 2013). We therefore also expect the effects on biotic properties to be more pronounced in a few years.

Another consideration is that deadwood affects soil microbial activity and biomass at a finer scale (Błońska et al., 2024; Minnich et al., 2021). Pooling the four soil samples from each patch before measuring soil biotic properties may dilute small-scale effects, such as those near deadwood. Given the high heterogeneity of soils (Lladó et al., 2018; Vos et al., 2013), short-term ESC effects might only be detectable at a small scale or could be overshadowed by other environmental factors.

Further, this study focused on only two broad soil functions (and their ratio, *i.e.,* respiratory quotient). We recommend future research to include testing for more specialized functions (*i.e.,* those functions carried out by a small group of specialized microorganisms), more precise indicators for functions (*i.e.*, measuring microbial carbon use efficiency using isotope labelling instead of the respiratory quotient), soil multifunctionality, and examining the community composition of soil organisms to understand their responses to enhanced structural complexity (Ali, 2023; Churchland et al., 2021; Lang et al., 2023; Lewandowski et al., 2015; Zheng et al., 2019).

### 4.3. Mechanistic context dependency of ESC effects (H3)

None of the measured abiotic soil variables was altered by ESC in the same way as the biotic variables; thus, the effects cannot simply be explained by a change in abiotic factors in the specific forest sites (*e.g.* nutrient availability increased in one forest site and decreased in another explaining the change of basal respiration). However, ESC induced changes in soil water content and understorey properties were linked to changes in microbial properties (H3). In forest sites with a higher increase in soil water content in response to ESC, soil microbial biomass increased more strongly. Notably, we did not find the same relationship for soil water content and soil basal respiration, even though soil water content is known to strongly affect soil basal respiration (Moghimian et al., 2020; Serna-Chavez et al., 2013; Siebert et al., 2023). Besides abiotic soil variables, changes in understorey vegetation might mediate ESC effects on soil microbial properties. However, no significant relationships were found between changes in understorey biomass or plant species richness and changes in microbial activity or biomass, respectively. This contradicts previous studies that have shown increased microbial activity and biomass with higher plant diversity in grasslands and forests (Chen et al., 2019; Eisenhauer et al., 2010, 2011; Lange et al., 2015; Strecker et al., 2016). While the effects might take longer to become apparent (Eisenhauer et al., 2010; Strecker et al., 2016), relationships in forests might differ from relationships in grasslands (Xu et al., 2020), and understorey plant species richness effects could be diminished by the tree canopy and the forest microclimate (De Frenne et al., 2021). Notably, a greater increase in understorey plant biomass led to a stronger decrease in the respiratory quotient, indicating higher substrate-use efficiency of the soil microbial community, as more microbial biomass can be built up with less respiration (Eisenhauer et al., 2013). A tendency towards the same effect was found for changes of understorey plant species richness on changes of the respiratory quotient. Possibly, these findings were the result of altered resource quality and quantity due to more root biomass and root exudates by the understorey plant community (Eisenhauer et al., 2017). However, when calculating the effect sizes for each forest site, we obtained only eight data points, so these results should be interpreted with caution. Nonetheless, these results highlight the potentially important role of soil water content and the understorey vegetation as key factors that connect enhanced structural complexity to increased soil microbial functioning. More detailed investigations of the microbial communities (e.g., taxonomic composition and functional diversity) may provide more insights.

To better understand the context-dependency of ESC effects, future studies should examine which site-specific factors and interactions influence responses to ESC. While increasing the sample size could help, this may be impractical; controlled experiments with varied environmental conditions and structural complexity offer a more feasible alternative for uncovering underlying mechanisms.

In addition to spatial variability, context-dependency may also occur over time. For example, deadwood effects on soil properties have been found to be stronger in summer than in autumn (Gonzalez-Polo et al., 2013). Since we sampled in autumn—when microbial diversity typically peaks (Du et al., 2018; Voříšková et al., 2014)—future studies should include multiple seasons to better capture temporal dynamics (Lewandowski et al., 2015, 2019).

### 4.4. Temporal development of ESC effects over the first five years (H4)

There was no significant effect of enhanced structural complexity in any of the years from 2019-2023 on soil basal respiration, microbial biomass, and soil respiratory quotient. Accordingly, we did not find an increasing effect over time (H4), suggesting that either the forest context does not favor an ESC effect or that it takes longer to materialize under these conditions. A former study also found no effect of canopy gaps on soil microbial biomass over the seven years after gap establishment, but found effects on the soil microbial community composition over the first four years (Lewandowski et al., 2015). This emphasizes the need to investigate microbial community composition in addition to the ecosystem functions. Findings about temporal effects from grassland studies, where the relationship between plant species richness and microbial properties strengthens over time (Strecker et al., 2016), may not directly apply to our setting. Forest gaps usually regenerate quickly under low ungulate browsing pressure, resulting in increased understorey biomass and plant diversity (Sabo et al., 2019). However, this natural regeneration process likely differs from how plant diversity develops in experimental grassland studies, where vegetation is established on bare soil. Nonetheless, the relationship between biodiversity and ecosystem functioning strengthening over time (Guerrero-Ramírez et al., 2017; Strecker et al., 2016) might still apply to forest understorey vegetation and soil microbial properties. We recommend investigating soil ecosystem functions over longer time frames in different forest sites to explore if treatment effects and their temporal development differ.

### 4.5. Conclusions

In conclusion, the focal soil abiotic and biotic properties showed remarkable resistance to interventions enhancing forest structural complexity. Despite these limited overall responses, the interventions had significant but variable effects on certain soil properties. For instance, while nutrient availability increased near deadwood, soil pH and water content remained unaffected – thus providing only partial support for H1. Contrary to H2, no general effect of enhanced structural complexity (ESC) on microbial activity, biomass, or the respiratory quotient was observed. Instead, responses varied between sites, highlighting the importance of site-specific conditions. Notable links between increased soil water content and microbial biomass, as well as between understorey plant biomass and the respiratory quotient (an indicator of microbial substrate use efficiency), support the context-dependent relationships proposed in H3. These findings indicate that the pathways connecting structural complexity and soil functioning are indirect, influenced by local conditions, and emphasize the important roles of soil moisture and understorey vegetation in shaping microbial communities. The anticipated temporal strengthening of effects (H4) was not supported within the conditions and timeframe of this study. To advance the development of practical forest management strategies that enhance soil ecosystem functioning, we encourage experiments that combine structural enhancement with active understorey management, controlled manipulation of deadwood decay stages, and assessments of microbial community composition and functional diversity, while covering longer timeframes. Such approaches will clarify how structural complexity can support both aboveground biodiversity and the soil processes essential for long-term forest resilience and ecosystem functioning.

## Acknowledgements

We thank Anja Zeuner, Aron Weiss, Anne Busch, Bennet Knienieder, Felix Zeh, David Moore, Alfred Lochner, Irina Kalmanova, Anne Peter, Rabea Klümpers, Clara Wild, Marina Wolz, Jens Schlüter, Josef Nußhardt, Magnus Kraatz, Anne Wendlandt, Clara Dembowski, and Christoph Stegen for their help in the field and laboratory. We also thank the entire BETA-FOR team for their support.

