## Supplementary material for "Inconsistent short-term effects of enhanced structural complexity on soil microbial properties across German forests": Supplemetary material

**Supporting information**

Table S1: F- / $\chi^2$ and p-values of ANOVA models for the effect of the spatial variant of tree removel (control vs. distributed vs. aggregated), canopy openness (proportion of visible sky), deadwood volume per patch ($m^3$), and deadwood occurrence at the sampling point on the soil water content, soil pH, soil carbon-to-nitrogen ratio (C:N ratio), soil C, and soil N. For data on patch level (soil water content), the forest site was used as a random effect. For data on subpatch level (soil pH, soil C:N ratio, soil C, and soil N), the patch nested in forest site was used as a random effect. For percentage data (soil water content, soil C, and soil N), beta regression models were fitted, and for non-percentage data (soil pH, soil C:N ratio), we used models with normally distributed errors. Significant factors are in bold. NA= factor not available for model.

| **Soil abiotic properties** | | **Soil water content [%]** | | **Soil pH** | | **Soil C:N ratio** | | **Soil C [%]** | | **Soil N [%]** | |
|---|---|---|---|---|---|---|---|---|---|---|---|
| **Factors** | Df | $\chi^2$ | p | F | p | F | p | $\chi^2$ | p | $\chi^2$ | p |
| **Spatial variant of tree removal** | 2 | 1.381 | 0.502 | 0.703 | 0.497 | 0.520 | 0.596 | 0.310 | 0.856 | 1.131 | 0.568 |
| **Canopy openness** | 1 | 0.740 | 0.390 | 2.302 | 0.132 | 1.284 | 0.259 | 0.000 | 0.999 | 0.254 | 0.614 |
| **Deadwood volume** | 1 | 0.280 | 0.597 | 0.469 | 0.495 | 0.534 | 0.466 | 0.015 | 0.904 | 0.147 | 0.701 |
| **Deadwood occurrence** | NA | NA | NA | 0.640 | 0.424 | **3.892** | **0.049** | **5.279** | **0.022** | **4.052** | **0.044** |

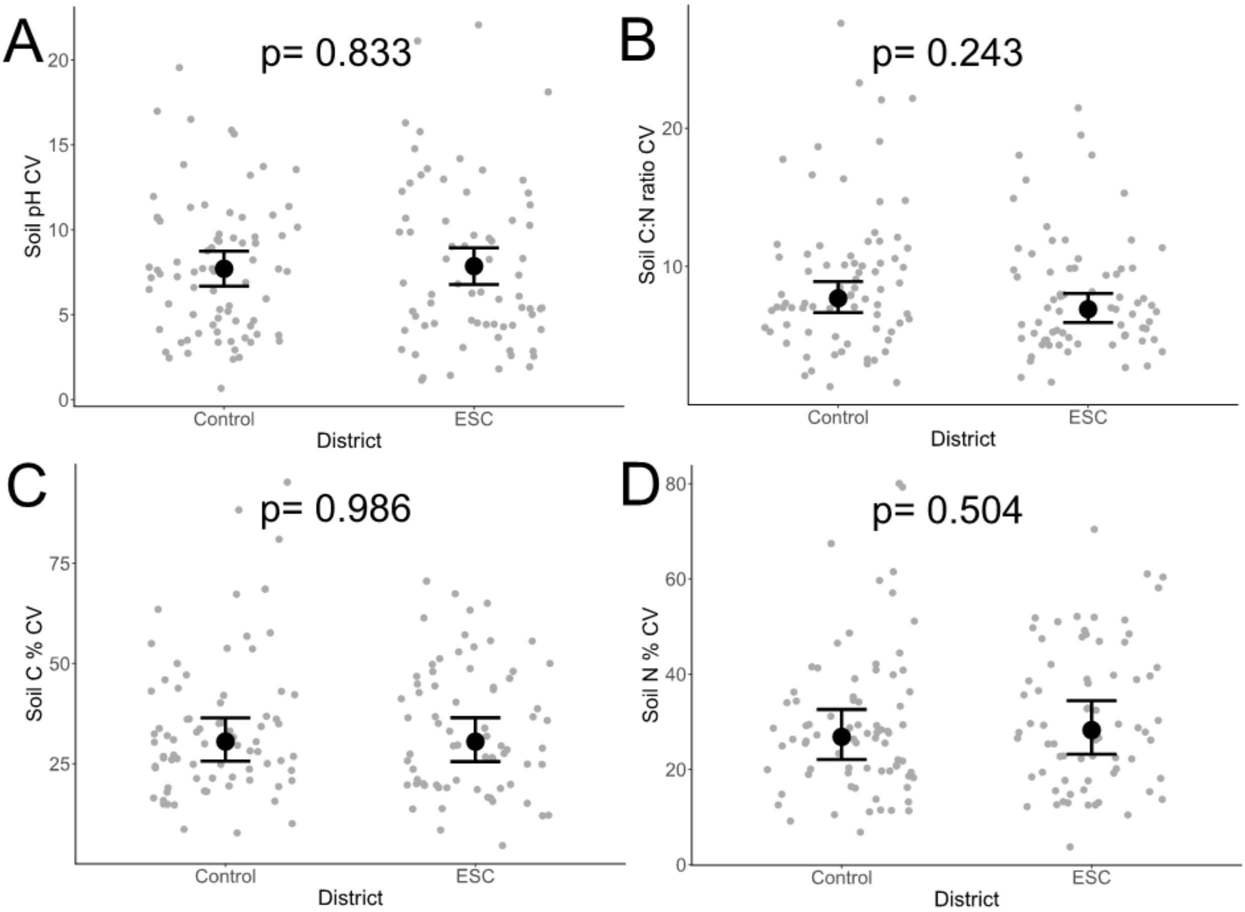


Figure S1: Coefficient of variation of soil pH (A), soil C:N (B), soil C% (C), and soil N% (D) in control and enhanced structural complexity (ESC) districts. P-values are derived from linear models. The model predictions are displayed with error bars representing the 95% confidence intervals and mean values indicated by larger points, while raw data points for each patch are shown in the background.

Table S2: F- and p-values of ANOVA models for the effect of district (enhancement of structural complexity vs. control) on the coefficient of variation (CV) of soil pH, soil carbon-to-nitrogen ratio (C:N ratio), soil C, and soil N. Forest site was used as a random effect.

| | **Soil pH CV** | | | **Soil C:N ratio CV** | | | **Soil C CV** | | | **Soil N CV** | | |
|---|---|---|---|---|---|---|---|---|---|---|---|---|
| **Factor** | Df | F | p | Df | F | P | Df | F | p | Df | F | p |
| **District** | 1 | 0.045 | 0.833 | 1 | 1.376 | 0.243 | 1 | 0.003 | 0.986 | 1 | 0.450 | 0.504 |

Table S3: Table of F- and p-values of ANOVA models for the effect of district (enhancement of structural complexity vs. control), forest site, and their interaction on soil basal respiration, soil microbial biomass, and the respiratory quotient for forest site U03. Significant factors are in bold.

| **Soil biotic properties** | | **Soil basal respiration** | | **Soil microbial biomass** | | **Respiratory quotient** | |
|---|---|---|---|---|---|---|---|
| **Factor** | Df | F | p | F | P | F | p |
| **District** | 1 | 1.779 | 0.185 | 3.048 | 0.083 | 0.249 | 0.619 |
| **Forest site** | 7 | **34.485** | **>0.0001** | **36.117** | **<0.0001** | **61.102** | **<0.0001** |
| **District: Forest Site** | 7 | **2.554** | **0.017** | 1.844 | 0.084 | 1.569 | 0.150 |

Table S4: Each row represents an individual linear model and shows the F- and p-values of the respective ANOVA model. We tested the correlations between the effect sizes of abiotic/understorey properties and the effect sizes of soil biotic properties. Significant factors are in bold. Effect sizes are the relative difference between enhanced structural complexity and control district for each property.

| | **Soil basal respiration effect size** | | | **Soil microbial biomass effect size** | | | **Respiratory quotient effect size** | | |
|---|---|---|---|---|---|---|---|---|---|
| **Factors** | Df | F | p | Df | F | p | Df | F | p |
| **Soil water content effect size** | 1 | 0.701 | 0.435 | 1 | **7.215** | **0.036** | 1 | 0.377 | 0.562 |

| | | | | | | | | | |
|---|---|---|---|---|---|---|---|---|---|
| **Soil C:N ratio effect size** | 1 | 0.669 | 0.444 | 1 | 0.603 | 0.467 | 1 | 0.014 | 0.910 |
| **Soil C% effect size** | 1 | 3.677 | 0.104 | 1 | 0.433 | 0.535 | 1 | 1.630 | 0.249 |
| **Understorey biomass effect size** | 1 | 0.440 | 0.502 | 1 | 0.510 | 0.532 | 1 | **7.324** | **0.035** |
| **Observed understorey species richness effect size** | 1 | 8.964 | 0.622 | 1 | 0.671 | 0.374 | 1 | 0.629 | 0.059 |

Table S5: Compilation of effect of district (D; control vs. enhanced structural complexity (ESC)) on soil properties per forest site (FS). Model output obtained from models (beta regression models for soil water content, soil C%, and soil N%, for the rest linear mixed-effect models) testing district, forest site and their interaction as explanatory variables (for basal respiration, microbial biomass, and the respiratory quotient, the device was additionally used as a random effect). Arrows show results of post-hoc tests in case of significant interaction of forest site and district. Yellow arrows represent an increasing effect of ESC and blue arrows a decreasing effect.

| | ANOVA model output | P08 | B05 | B07 | L11 | B06 | B04 | S10 | U03 |
|---|---|---|---|---|---|---|---|---|---|
| **Basal respiration** | D p=0.185<br>FS p<**0.0001**<br>D:FS p=**0.017** | ↓ | ns | ns | ↑ | ns | ns | ns | ns |
| **Microbial biomass** | D p=0.083<br>FS p<**0.0001**<br>D:FS p=0.084 | ns | ns | ns | ns | ↑ | ns | ns | ns |
| **Respiratory quotient** | D p=0.619<br>FS p<**0.0001**<br>D:FS p=0.150 | ns | ns | ns | ns | ns | ns | ns | ns |
| **pH** | D p=0.128<br>FS p<**0.0001**<br>D:FS p=0.260 | ns | ns | ns | ns | ns | ns | ns | ns |
| **Water content** | D p=**0.009**<br>FS p<**0.0001**<br>D:FS p<**0.0001** | ns | ns | ns | ↑ | ↑ | ns | ↑ | ns |
| **C:N** | D p=0.811<br>FS p<**0.0001**<br>D:FS p=**0.013** | ↓ | ns | ns | ns | ns | ↑ | ns | ns |
| **C %** | D p=0.195<br>FS p<**0.0001**<br>D:FS p=0.052 | ns | ↓ | ns | ns | ns | ns | ns | ns |
| **N %** | D p=0.127<br>FS p<**0.0001**<br>D:FS p=**0.016** | ns | ↓ | ns | ns | ns | ns | ns | ns |

Table S6: F- and p-values of ANOVA models for the effect of district (enhancement of structural complexity vs. control), year, and their interaction on soil basal respiration, soil microbial biomass and the respiratory quotient for forest site U03. Patch and device were used as random effects. Significant factors are in bold.

| | **Soil basal respiration** | | | **Soil microbial biomass** | | | **Respiratory quotient** | | |
|---|---|---|---|---|---|---|---|---|---|
| **Factor** | Df | F | p | Df | F | p | Df | F | p |
| **District** | 1 | 0.054 | 0.816 | 1 | 0.549 | 0.461 | 1 | 0.471 | 0.494 |
| **Year** | 1 | **26.640** | **<0.0001** | 1 | 0.040 | 0.842 | 1 | **49.455** | **<0.0001** |

| | | | | | | | | | |
|---|---|---|---|---|---|---|---|---|---|
| **District: Year** | 1 | 0.054 | 0.816 | 1 | 0.547 | 0.461 | 1 | 0.470 | 0.494 |

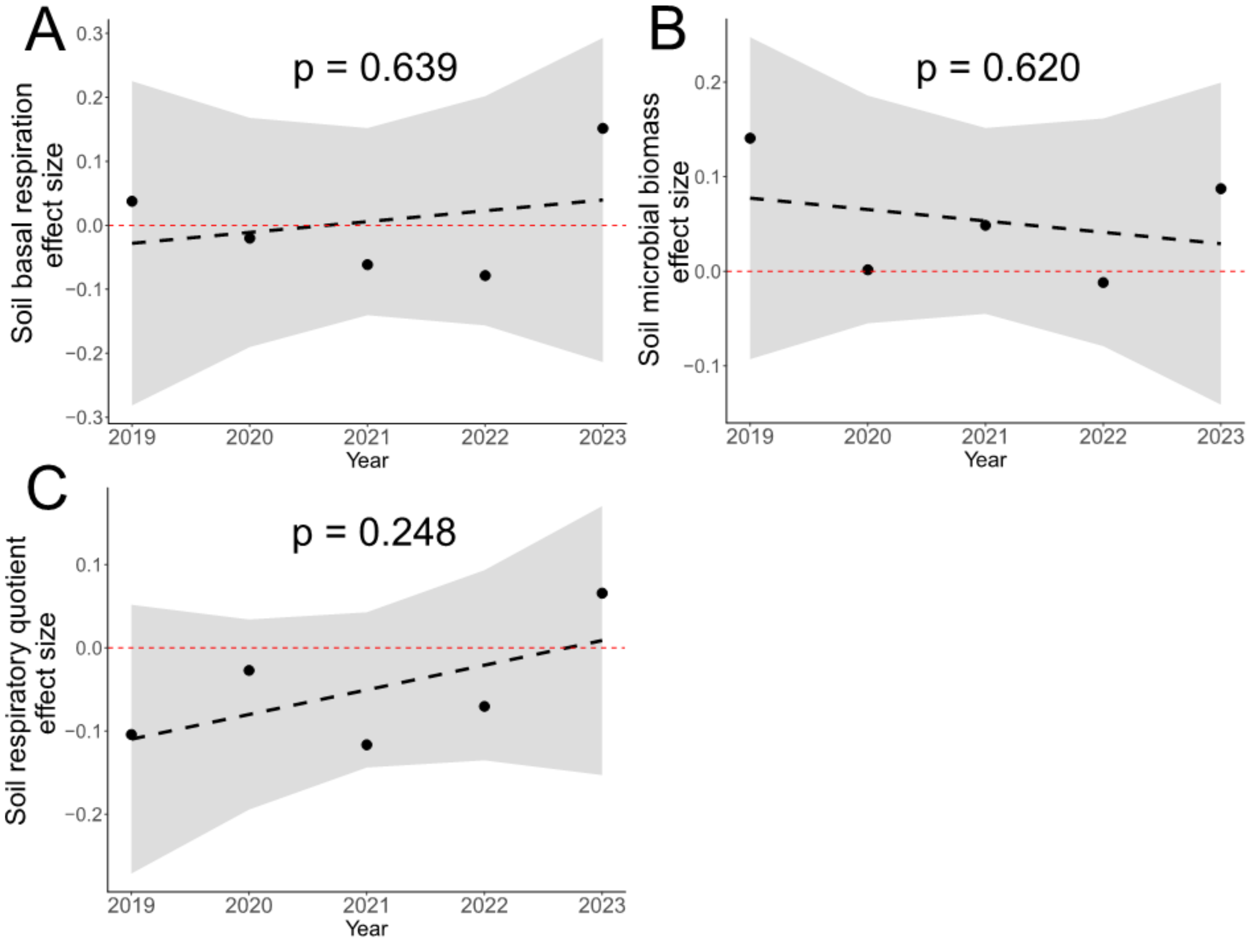


Figure S2: The effect size (relative difference between enhanced structural complexity (ESC) and control district) for soil basal respiration (A), soil microbial biomass (B), and respiratory quotient (C) for the years 2019-2023 in the forest site U03. P-values are derived from linear models. The black line represents, respectively, the predicted values with the 95% confidence interval, the points show the raw data. Non-significant relationships are displayed with dashed black lines. The red line is the zero point on the y-axis and therefore values above it show an increasing effect and values below it a decreasing effect by ESC.

Table S7: F- and p-values of ANOVA models for the effect of the year on the the effect size (relative difference between enhanced structural complexity and control district) for soil basal respiration, soil microbial biomass, and respiratory quotient in the forest site U03.

| | **Soil basal respiration effect size** | | | **Soil microbial effect size** | | | **Respiratory quotient effect size** | | |
|---|---|---|---|---|---|---|---|---|---|
| **Factor** | Df | F | p | Df | F | p | Df | F | p |
| **Year** | 1 | 0.271 | 0.639 | 1 | 0.304 | 0.620 | 1 | 2.043 | 0.248 |
